# Diagnostically distinct resting state fMRI energy distributions: A subject-specific maximum entropy modeling study

**DOI:** 10.1101/2024.01.23.576937

**Authors:** Nicholas Theis, Jyotika Bahuguna, Jonathan E. Rubin, Joshua Cape, Satish Iyengar, Konasale M. Prasad

## Abstract

**Objective:** Existing neuroimaging studies of psychotic and mood disorders have reported brain activation differences (first-order properties) and altered pairwise correlation-based functional connectivity (second-order properties). However, both approaches have certain limitations that can be overcome by integrating them in a pairwise maximum entropy model (MEM) that better represents a comprehensive picture of fMRI signal patterns and provides a system-wide summary measure called energy. This study examines the applicability of individual-level MEM for psychiatry and identifies image-derived model coefficients related to model parameters.

**Method:** MEMs are fit to resting state fMRI data from each individual with schizophrenia/schizoaffective disorder, bipolar disorder, and major depression (n=132) and demographically matched healthy controls (n=132) from the UK Biobank to different subsets of the default mode network (DMN) regions.

**Results:** The model satisfactorily explained observed brain energy state occurrence probabilities across all participants, and model parameters were significantly correlated with image-derived coefficients for all groups. Within clinical groups, averaged energy level distributions were higher in schizophrenia/schizoaffective disorder but lower in bipolar disorder compared to controls for both bilateral and unilateral DMN. Major depression energy distributions were higher compared to controls only in the right hemisphere DMN.

**Conclusions:** Diagnostically distinct energy states suggest that probability distributions of temporal changes in synchronously active nodes may underlie each diagnostic entity. Subject-specific MEMs allow for factoring in the individual variations compared to traditional group-level inferences, offering an improved measure of biologically meaningful correlates of brain activity that may have potential clinical utility.

## Introduction

Neuroimaging studies of schizophrenia and mood disorders have revealed important findings that merit investigation using more innovative and integrative methods to advance our understanding of pathophysiology and develop novel treatments. Existing approaches to find group differences in regional activations (first-order properties) do not consider how activation difference in one region may influence differences in another region, and methods relying on correlation-based coactivations of node pairs (second-order properties, e.g., functional connectivity, FC) do not reflect the magnitudes of regional activations and are often not time-resolved. These limitations can be overcome by mathematically reducing both the first- and the second-order properties to a scalar quantity called energy using the generalized Ising model, also called the maximum entropy model (MEM).

This model can describe elements of neural physiology (1–3) and characterize the probability that a brain network will be in any observable state representing a combination of regional activations and interactions. Multiple cortical areas, subcortical nuclei, and their complex connectivity engender whole-brain dynamics (4, 5) that can be assessed using blood oxygenation level-dependent (BOLD) signals (6). Temporal snapshots of this time-evolving system can be considered states in the same sense used in the MEM, though these states are binary simplifications of the underlying biological activity. Previous studies have observed that the interaction matrix of the MEM is related to traditional correlation-based FC but provides a more accurate representation of the underlying biology (7, 8). The physiological relevance of the model affords the opportunity to better understand the changes to brain connectivity associated with disorders such as schizophrenia, bipolar disorder, and major depression (9).

Earlier studies on healthy individuals have shown that concatenated functional magnetic resonance imaging (fMRI) state occurrence patterns across individuals can be captured using MEM (10–13), but only a few (7, 8, 14) have applied it to psychiatric disorders (15, 16). Most existing studies of the MEM have used concatenated data across multiple participants to produce a single, group-level model that cannot be applied to individuals without the possibility of ecological fallacy of using group level means for individual participants (17). Concatenation is done due to the need for many functional volumes to fit systems of even moderate size. If unequal state probabilities are expected, then the rarest states may be undetectable unless many more state observations are made than possible unique states. The number of possible unique states in a system where regional activity is binary (“on” or “off”), is given by 2*^N^*, where *N* is the number of regions. For 10 nodes, the number of unique states would be 1,024, which is larger than the number of volumes acquired in the typical fMRI (18).

Recent developments, relating to faster (with shorter repetition time, TR) and/or longer imaging sequences and algorithmic improvements have made it possible to achieve individual level MEM fitting (7, 16), which offers potential for clinical applications in psychiatry. Here, we focus on the development of image-derived coefficients for individuals that can help distinguish apparently divergent pathologies underlying different disorders. Potential clinical implications of FC (19, 20) could be tested more rigorously in specific systems with MEM.

We performed MEM fitting on regions included in the default mode network (DMN) in a transdiagnostic sample (N=132) and demographically matched controls (N=132) from the UK Biobank (21). The multimodal Glasser cortical parcellation was used to identify DMN regions (22) using published data (23). Alterations in DMN connectivity during resting state have long been associated with schizophrenia, depression, and other mood disorders (24–28). Our hypotheses were that 1) the MEM would fit data from various subsets of regions from the DMN parcels selected from the Glasser atlas in each individual, 2) the parameters of the MEM would relate to measures derived from traditional FC, and 3) state energies given by the model, which are inversely related to state probabilities, would differ by each diagnostic group compared to non-psychiatric controls.

## Methods

### Data Source

Imaging data was sourced from the UK Biobank (www.ukbiobank.ac.uk), a biomedical database of more than 500,000 participants (21), including MRI data for some participants (29). The following UK Biobank data fields were used to obtain psychiatric disorders defined according to the International Classification of Disease, 10^th^ edition: 130874 Schizophrenia (F20); 130884 Schizoaffective disorder (F25); 130890 Manic Episode (F30); 130892 Bipolar Affective Disorder (F31); and 130896 Recurrent Depressive Disorder (F33).These were combined into three groups, namely schizophrenia/schizoaffective disorder (n=17), bipolar disorder (n=40) and major depressive disorder (n=75) (total N=132).

We included 132 controls from the UK Biobank individually matched for age and sex to each ICD disorder-diagnosed individual. Controls were screened by excluding persons with the above fields as well as the field 20461, “age when first had unusual or psychotic experience”. Only subjects who had T_1_-weighted image and a resting fMRI data were included in the study.

### Imaging methods

Image acquisition parameters in the UK Biobank are published (30). Briefly, all UK Biobank sites used identical Siemens Skyra 3T scanners with standard Siemens 32-channel receiving coils. T_1_w images were acquired with a 3D MPRAGE sequence, 1mm isotropic voxels. For fMRI, a gradient echo-EPI sequence was used with x8 multi-slice acceleration, 2.4mm isotropic voxels, and a TR of 0.735 seconds for 490 timepoints. The imaging data were internally quality controlled and preprocessed (30). For fMRI, preprocessing steps included motion correction with MCFLIRT (31), grand-mean intensity normalization for the entire 4D acquisition, high-pass temporal filtering, EPI unwarping, gradient distortion correction, and ICA+FIX artifact removal (32, 33).

Additional image processing was performed using FSL and Freesurfer on the Pittsburgh Supercomputing Center’s Bridges-2 system (34). These steps included Human Connectome Project (HCP) atlas parcellation (22) using the Neurolab (35) approach where an fs_average version of the HCP multi-modal atlas parcellation (36) was used to register the atlas to the individual result of FreeSurfer’s “recon-all” function (37). Image co-registration of atlas parcellations to the fMRI was performed with FSL (31). MATLAB was used for voxel-wise averaging to generate nodal timeseries for each HCP parcel.

### Region Selection

The DMN consisted of 17 HCP regions per hemisphere (23), namely 10r, 31a, 31pd, 31pv, a24, d23ab, IP1, p32, POS1, POS2, RSC, PFm, PGi, PGs, s32, TPOJ3, and v23ab. A system of 34 nodes is beyond practical size for the MEM. We selected anatomically diverse nodes to create three subsets composed of size ≤ 9 regions. An 8-node bilateral system consisting of 4 symmetrical regions from each hemisphere and two 9-node unilateral systems that are mirror images of each other were selected. For the bilateral system, we selected POS1 (parietal-occipital sulcus), p32 (anterior cingulate cortex), TPOJ3 (temporo-parieto-occipital junction 3), and PGi (inferior parietal cortex) from each hemisphere. Each unilateral system consisted of anterior cingulate cortex (10r, a24, S32), posterior cingulate cortex (31pv, d23ab), area intraparietal (IP1), retrosplenial cortex (RSC), inferior parietal cortex (PFm), and TPOJ3 that includes the three anatomically diverse components included in the bilateral system. For a visual representation of these regions see **Supplemental Figure S1.**

### The generalized Ising Model

The generalized Ising model, often used synonymously with pairwise MEM (38), is “generalized” in the sense that all node-pairs interact directly, whereas the originally proposed model is more limited (39, 40). The pairwise MEM considers up to the second-order interaction (41). The model describes a system where the probabilities of occurrence of nodal binary on/off configurations (called states) are determined by the interactions of the elements of the system in a network responding to an external field (42). According to the model, states for any subset of brain regions-of-interest will exhibit an energy given by the generalized Ising equation:

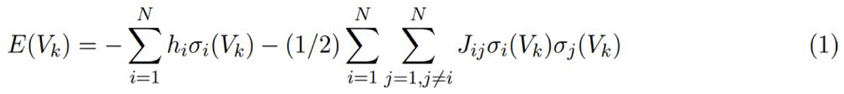

where the term energy, *E*, is defined for each brain state, *V_k_*. This equation defines the energy of a given state, *E(V_k_)*, as the sum of first-order activation terms and half the sum of all second-order co-activations (11). The size of the system (*N*) is the number of nodes on which data are collected. Each *σ_i_* term is binary, such that each *σ_i_* ∈ {0,1}, and indicates whether, within state *V_k_,* the activation of node *i* is above or below a threshold. For a given state number, *k*, the state *V_k_* is therefore a collection of *N* binary values representing binarized BOLD intensity at the imaged brain regions, and there are *2^N^* unique system states. Each term *h_i_* in equation (1) is the importance of the activation of the *i*^th^ node in the sense that certain nodes may have a greater influence over system dynamics than others. Each term *J_ij_* encodes the relative importance of the co-activation of a pair of regions.

The energy term, *E*, can be thought of as an abstract form of potential energy with no absolute value or physical units. The energy of a given state is related to the negative log probability of that state, *P(V_k_)*, according to the Boltzmann distribution:

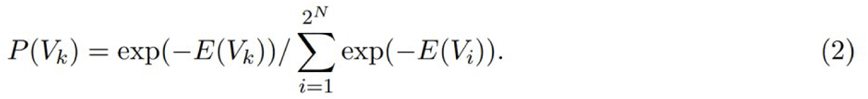

Using a computational approach to map from observed probabilities, *P(V_k_)* to unknown Ising model parameters, *J* and *h*, is called inverse Ising inference, and was performed using the maximum likelihood estimation (MLE) (7, 16) (see below).

### Observed Binary States (*B*)

After regional voxel averaging, fMRI signals are *z*-scored per subject per node. *Z*-scoring is widely used in fMRI studies (43) since the units of the original BOLD signal are arbitrary and have unknown baseline offsets. Next, the global signal across all nodes is subtracted out, which is similar to global signal normalization (44). Time series processing steps for a single subject is shown in **Figure 1**.

**Figure 1.**
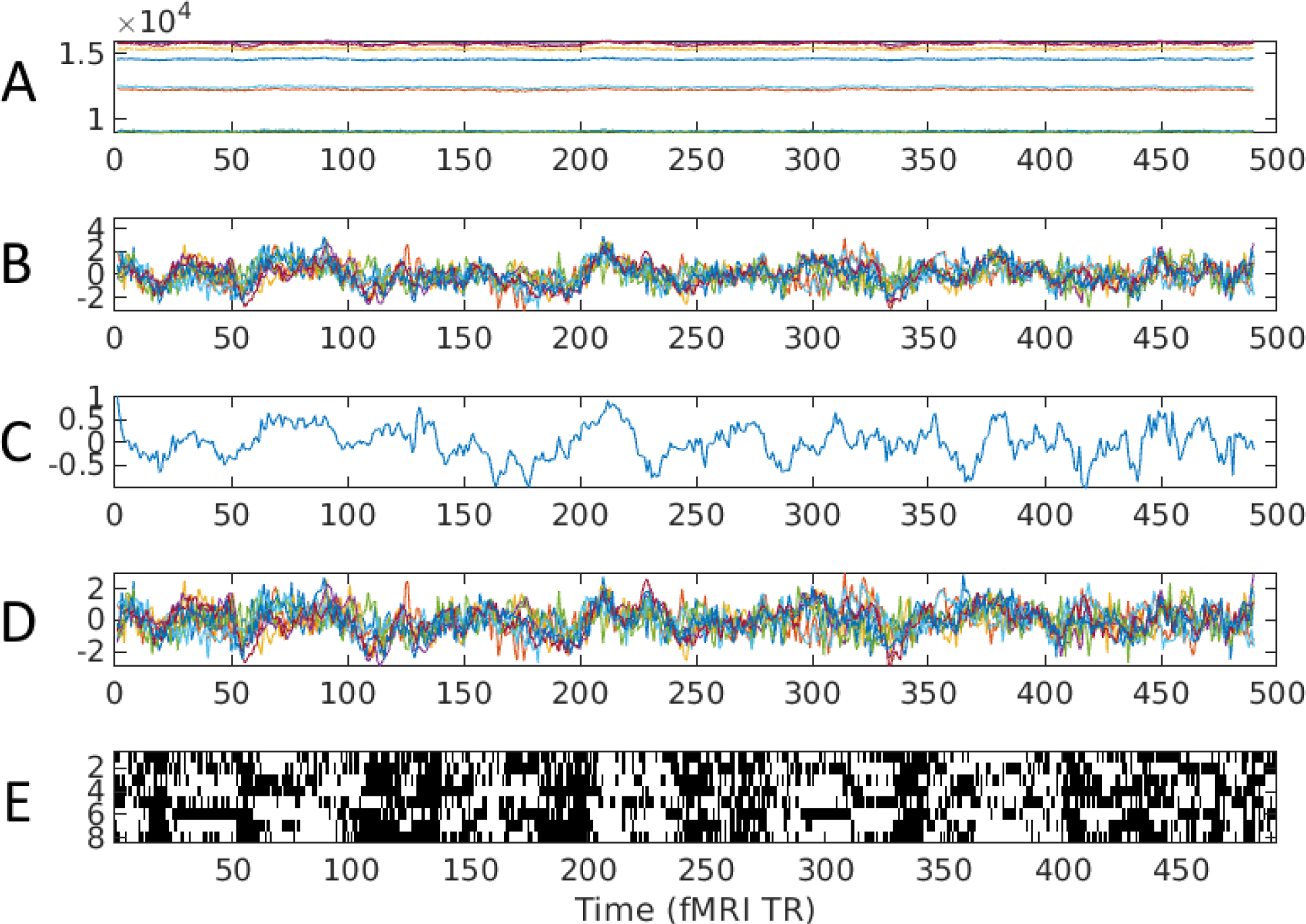
Post-processing and Binarization. All x-axes show time in fMRI volumes, Repetition Time (TR); each TR is 0.735 seconds. **A)** The preprocessed BOLD signals in amplitude of arbitrary units for a single subject for the 8 bilateral nodes: POS1, p32m aTPOJ3, and PGi in both hemispheres. **B)** The z-scored nodal time-series has an average centered at zero and is scaled to a standard deviation of one. **C)** The global signal across all z-scored nodes in the entire brain is shown, followed by **D)** the z-scores after subtracting this value. Finally, **E)** the time series are binarized at the threshold of zero, where the y-axis is now node index, rather than amplitude. White cells indicate the “on” nodes at the given time, meaning that the value in **D** for that node was greater than zero, while black cells indicate “off” nodes.

To generate a binary matrix *B* representing brain states from regional time series data, a threshold of zero is applied to the data for each region at each time point, with positive values set to 1 and negative values to 0. The resulting values are assembled in a matrix, *B*, where each row represents a region, and each column represents a time. The mean activation level for each region is by definition zero after z-scoring resulting in a matrix *B* with 50% “on” states and 50% “off” states (i.e., 50% of entries in each row of matrix *B* are 1 and 50% are 0). This threshold, zero, has previously been determined to be an optimal choice in terms of fit of MEM to empirical observations (15). Then, to calculate the empirical probability of state occurrence, for each subject, for each unique state the number of times that state appears during that subject’s acquisition is divided by the number of time samples, in this case 490.

### Maximum Likelihood Estimation (MLE) of the MEM

The maximum-entropy estimate was first proposed by Edwin Jaynes to find “maximally noncommittal” solutions to statistical inference in systems with missing information (45). The principle of maximum entropy provides a robust optimization criterion, but specific optimal parameters for the MEM must be obtained computationally. One approach to finding the coefficients of the MEM is applying gradient descent to perform MLE, based on the log-likelihood function given the set of observed brain states (7, 16). The MLE is an iterative method that continues to iterate on parameter estimates until a convergence criterion is met. While improvements on the method exist, e.g., the variational expectation-maximization algorithm, the MLE is well suited for our purposes because it does not require group level aggregates.

### Statistical Comparisons

A summary of model fit was evaluated after convergence by correlating the predicted probability given by calculating *P(V_k_)* for each state in *V_k_* using equations (1) and (2) to the corresponding observed probability for each subject’s binary state sequence. The strength of this correlation indicates the validity of the model. A two-sample Kolmogorov-Smirnov test by group is used to compare fitness distributions for group-level differences in each model. Additionally, edgewise coefficients from *J* are compared to FC, which is given by Pearson’s correlation (*r*) between node-pair timeseries before binarization, and node-wise coefficients from *h* are compared to the negative nodal strengths derived from FC, where the strength of a node is defined as the sum of entries in the row *or* column to which the node belongs (measure of how strongly the node covaries with other nodes of the network) in the matrix of network FC values:

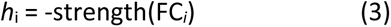

The relationship of the model parameters *J* and *h* to the estimates given by FC and equation (3), respectively, were tested using Pearson correlation either across all edges in the case of *J* or across all nodes in the case of *h*. Finally, the energy profiles, *E(V_k_)*, for each state *V_k_* were averaged across all subjects in each clinical group and across the appropriate control group. Each patient subgroup had a unique subset of control subjects which were individually demographically matched. The resulting group-averaged energy distributions were then compared using a two-sample Kolmogorov-Smirnov test for differences in distributions and visualized as histograms.

## Results

### Demographics

The groups did not differ significantly for age and sex since we selected control subjects matched for every subject with an ICD-10 diagnosis (**Table 1)**.

**Table 1.** Demographic data. Schizophrenia (SZ) and schizoaffective disorder (SA) are considered together to select controls to match for age and sex.

|  | Schizophrenia/<br>Schizoaffective | SZ/SA<br>Controls | Bipolar/<br>Manic | B/M<br>Controls | Depression | Depression<br>Controls |
| --- | --- | --- | --- | --- | --- | --- |
| Sample Size | 17 | 17 | 40 | 40 | 75 | 75 |
| Age (in<br>years) | 52.6 ($\pm 6.5$) | 52.6 ($\pm 6.5$) | 53.4 ($\pm 7.3$) | 53.4 ($\pm 7.3$) | 53.6 ($\pm 7.4$) | 53.6 ($\pm 7.4$) |
| % Female | 70.6% | 70.6% | 55.0% | 55.0% | 72.0% | 72.0% |

### Model Fit

Individual subject MLE fits of the MEM, evaluated as the correlation between the observed and predicted state occurrence probabilities, were statistically significant in all subjects in both groups and in all subsets of nodes examined (**Figure 2**). Most correlations (787/792) exceeded 0.9 (average *r*=0.91±0.064; p<1e-62) while the rest of models had correlations less than 0.5 but were nonetheless significant (p<0.1e-23). In contrast, using FC directly to determine the values of *J* and equation (3) for *h* resulted in weaker correlations between observed and predicted states, where no correlation exceeded 0.7 (**Supplemental Figure S2**). Group differences were not found in the model fits. Specifically, in all three systems, a KS-test found no significant differences between the distributions of model fitness in patients versus controls (bilateral: KS=0.152, p=0.087; left: KS=0.114, p=0.342; right: KS=0.091, p=0.626).

**Figure 2.**
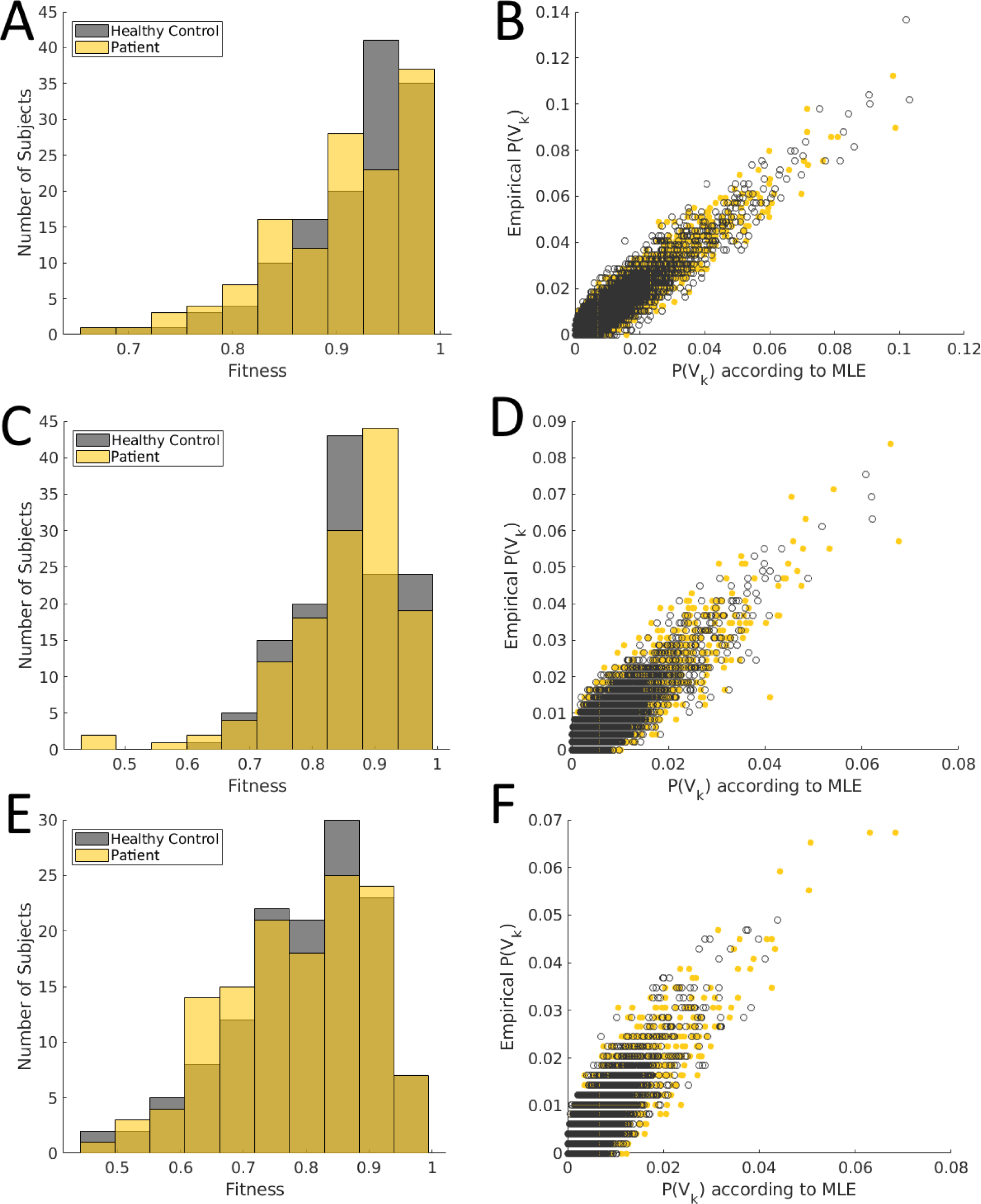
**MEM Fits. Bilateral DMN nodes** (top row) **A:** Two histograms, psychiatric disorders group (N=132) in yellow and pooled HC (N=132) in black, are compared and not significantly different by a KS-test (KS=0.152, p=0.087). Fitness (x-axis) is the correlation between predicted P(V_k_) according to the model and observed P(V_k_) per individual. **B)** All unique possible states are shown for all subjects according to their probability predicted from the MEM (x-axis) versus their observed probability (y-axis). Black circles represent data from controls, yellow points represent data from the patient group. **Left Hemisphere DMN nodes (**Middle row) **C:** Fitness histograms for pooled patient group (yellow) and pooled HC (black) are compared and not significantly different (KS=0.114, p=0.342). **D)** All unique possible states are shown for all subjects according to their predicted probability (x-axis) versus their observed probability (y-axis). Black circles represent data from controls, yellow points represent data from the patient group. **Right Hemisphere DMN nodes** (Bottom row) **E:** Fit histograms for pooled patients (yellow) and pooled HC (black) are compared and not significantly different (KS=0.091, p=0.626). **F:** All unique possible states are shown for all subjects according to their predicted probability (x-axis) versus their observed probability (y-axis). Black circles represent data from controls, yellow points represent data from the patient group.

### Similarity of MEM Parameters to FC

There were significant correlations across participants between MEM parameters and image-derived coefficients in all individuals and subsets of nodes (**Figure 3**). The distributions of these correlations were not significantly different between subjects with all psychiatric disorders, taken as a group, and controls. The relationship between *J* and FC appears to be sigmoidal with a linear relation outside of the saturating part of the sigmoid. The nature of this relationship is explained by the fact that FC is bounded by 1 but the entries of *J* are not bounded. The correlation between *h* and the image-derived coefficients associated with *h* at the individual level, computed from equation (3), was stronger than the *J*-FC correlation in most subjects.

**Figure 3.**
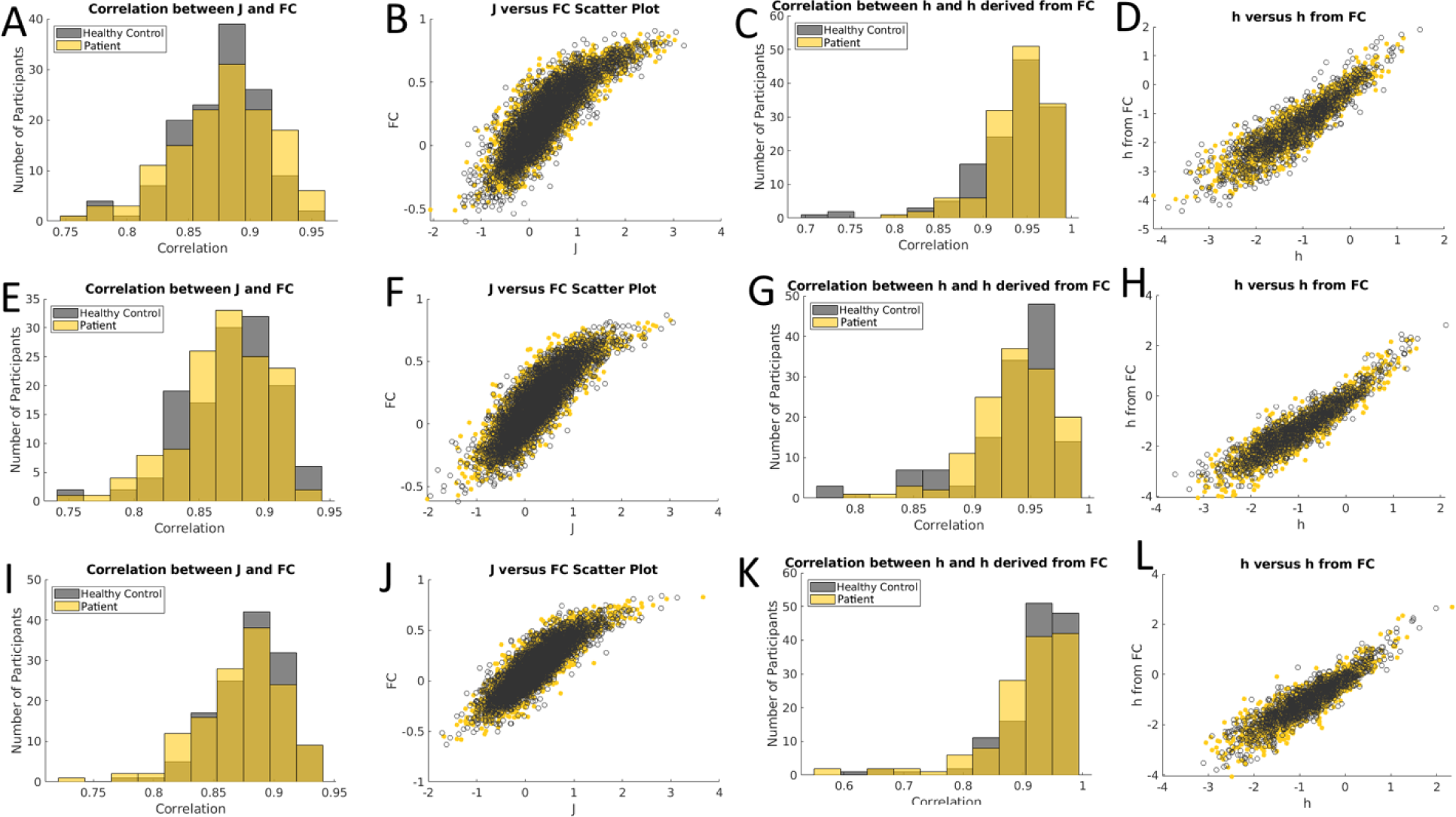
Comparison of MLE- and image-derived MEM coefficients. **<u>Bilateral</u> DMN nodes** (top row). **A)** Two histograms, pooled patients (N=132) in yellow and pooled HC (N=132) in black are compared. Each histogram represents the population-wide distribution of correlations between the second-order J parameter (x-axis) compared to edge weights of the FC (y-axis). **B)** Scatter plots of all subjects and all parameter values showing a strong relationship between J and FC; black circles are HC; yellow points are patients. **C)** Histograms for group-wise distributions of correlations between h from the MEM and the image-derived coefficients in equation (3). **D)** Individual scatter plots for all participants between h (x-axis) and the image-derived coefficients from equation (3) (y-axis). **Left Hemisphere DMN nodes** (Middle row) **E)** Group-wise population histograms of J-to-FC similarity. **F)** All individual-level scatter plot data for corresponding parameters in J (x-axis), and FC (y-axis). **G)** Histograms for group-wise distributions of correlations between h from the MEM and the image-derived coefficients from equation (3). **H)** Individual scatter plots for all participants between h (x-axis) and the image-derived coefficients (y-axis). **Right Hemisphere DMN nodes (**Bottom row) **I)** Group-wise population histograms of J-to-FC similarity. **J)** All individual-level scatter plot data for corresponding parameters in J (x-axis), and FC (y-axis). **K)** Histograms for group-wise distributions of correlations between h from the MEM and the image-derived coefficients from equation (3). **H)** Individual scatter plots for all participants between h (x-axis) and the image-derived coefficients (y-axis).

### Energy Profiles by Clinical Group

In the bilateral DMN system, both the bipolar disorder (KS=0.406, p=3.38e-19) and the SZ/SZA energy distributions were significantly different from controls (KS=0.301, p=1.025e-10) (**Figure 4A-C**) but the energy distributions across states for the depression group and the controls did not differ (KS=0.090, p=0.241). In the left hemisphere DMN system, the group-wide average energy profiles followed the same trends as the bilateral case for the SZ/SZA, bipolar disorder and depression groups (**Figures 4D-F**). The right hemisphere DMN system followed a similar trend as the bilateral system with bipolar disorder (KS=0.315, p=9.20e-23) and SZ/SZA (KS=0.404, p=2.38e-37) having significantly different distributions; unlike the bilateral and left hemisphere DMN systems, the depression group in this case was found to have a significantly different energy distribution from the demographically matched HC (KS=0.127, p=4.57e-04) (**Figure 4G-I**). Specifically, the energy state distributions for schizophrenia/schizoaffective were in the higher range compared to controls in all systems of DMN considered here. However, bipolar disorder patients had average energy state distributions that were significantly lower than controls.

**Figure 4.**
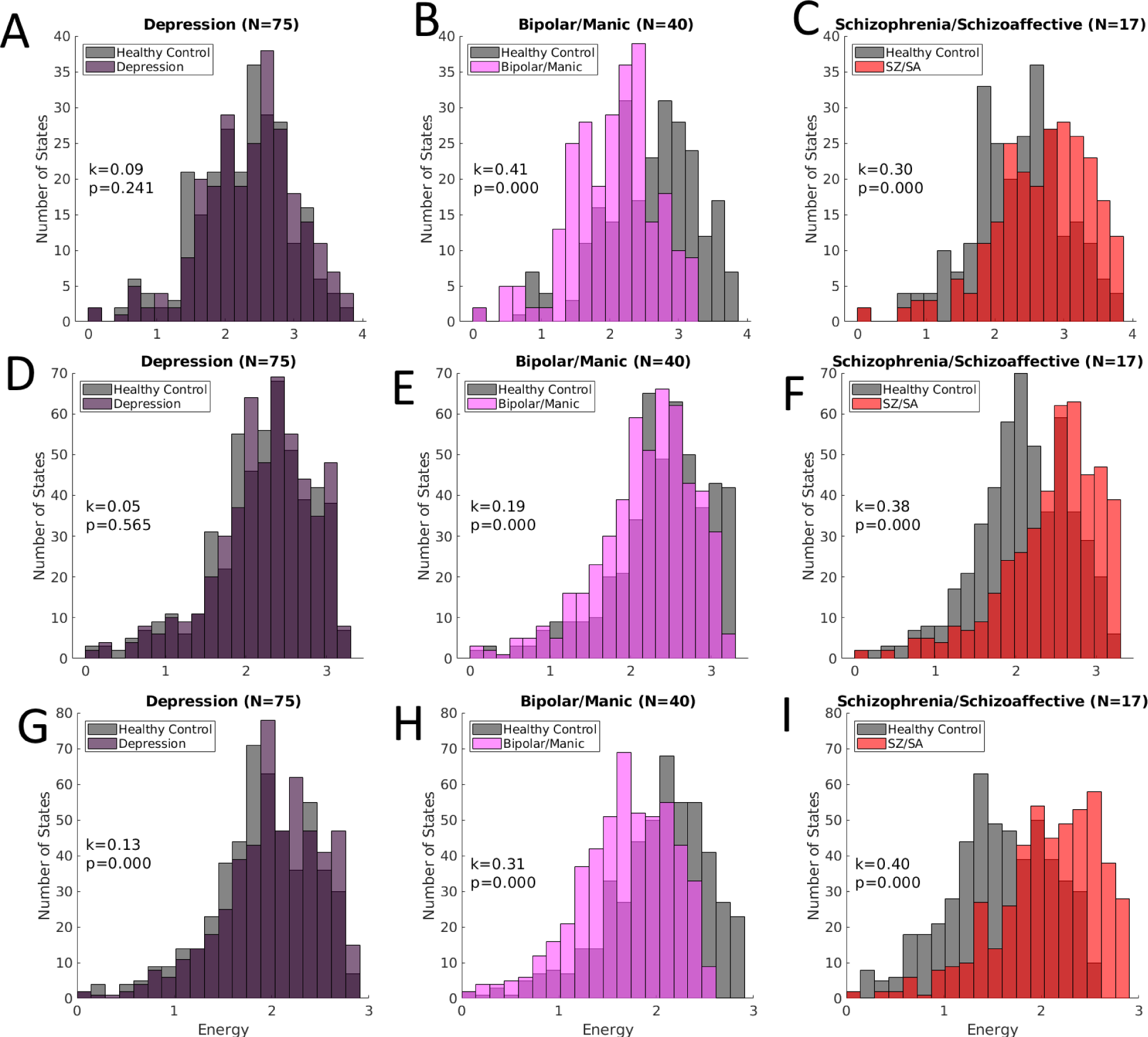
Energy Profiles by Clinical Group. **Bilateral:** top row. **Left Hemisphere:** middle row. **Right Hemisphere:** bottom row. Statistics reported in plots are KS statistics and associated p-value. **A) bilateral:** The group-wise average energy profile of depression compared to controls. **B) bilateral:** bipolar/manic compared to controls. **C) bilateral:** SZ/SA compared to controls. **D) left hemisphere:** depression compared to controls. **E) left hemisphere:** bipolar/manic compared to controls. **F) left hemisphere:** SZ/SA compared to controls. **G) right hemisphere:** persons with depression compared to controls. **H) right hemisphere:** bipolar/manic compared to controls. **I) right hemisphere:** SZ/SA compared to controls.

#### *Post-hoc* comparisons among the clinical groups

The energy distribution among schizophrenia/schizoaffective disorder was significantly different from that of bipolar energy distribution in all three systems (bilateral: KS=0.461, p=1.36e-24; left: KS=0.307, p=1.15e-21, right: KS=0.357, p=2.77e-29), with the schizophrenia/schizoaffective group’s energy distribution shifted towards higher values in all systems. Similarly, schizophrenia/schizoaffective disorder group showed significant differences compared to the depression group in all three systems (bilateral: KS=0.281, p=2.01e-09; left: KS=0.285, p=8.56e-19; right KS=0.154, p=6.43e-30), with the depression group’s energy distributions shifted towards lower values. However, the bipolar group and the depression group only had significant differences in energy distributions in the bilateral system (KS=0.266, p=1.89e-08) and the right hemisphere system (KS=0.305, p=2.14e-21) with the bipolar group associated with lower energies; differences between these groups were not in the left hemisphere (KS=0.043, p=0.723).

## Discussion

Our main finding is that the DMN abnormalities of connectivity and intrinsic activity is detectable and may be diagnostically distinct using the MEM framework that obtains an integrative and novel measure of neural dynamics. Further, the MLE algorithm for inferring MEM coefficients converged across all node collections examined in every subject (7, 16) regardless of the clinical status, and the correlation between the predicted and the observed *P(V_k_)* distributions for each individual was significant (**Figures 2**) and exceeded the correlation derived from FC (**Supplemental Figure S**) suggesting that MLE fit for MEM provides a robust method for examining the energy of states among each individual. Moreover, the image-derived coefficients from FC correlate with the model parameters *h* and *J* (**Figure 3**), suggesting that the MEM could be an alternative estimate of biological activation that better represents overall brain state patterns.

To our knowledge, we are the first to observe diagnostically distinct distributions of brain state energy among the three ICD-10-defined clinical groups. However, correlation with clinical measures could not be done because fine-grained psychopathological and cognitive measures were not available. However, from a neurobiological perspective, a prior study on schizophrenia, major depression, and bipolar disorder compared to controls did not find DMN dysconnectivity as a transdiagnostic feature. However, DMN dysconnectivity was more specific to schizophrenia (46), which was similar to the findings in a metanalysis (47). The DMN regions with dysconnectivity in bipolar disorder and major depression were distinct compared to DMN regions with dysconnectivity in schizophrenia (46). A study using *dynamic* functional connectivity on a similar transdiagnostic group reported that the three disorders exhibited similar dynamic functional dysconnectivity except that schizophrenia showed more severe changes compared to bipolar disorder and depression (48). Since the MEM estimates activation weights (*h*) and interaction strengths (*J*) from system-wide patterns, we consider it to be statistically more robust than traditional pairwise correlation-based connectivity models. MEM approach provides a temporal resolution at each TR accounting for non-stationarity of brain activity and connectivity much better than dynamic functional connectivity that combines multiple TRs to infer the temporal changes. Additionally, the generalized Ising model **Equation 1** combines these first-order and second-order parameters into a single *probability model* (**Equation 2**) that better estimates the observed brain state likelihoods.

Differences across the disorders in the energy values associated with the states of a specific system (**Figure 4**) may also have pathophysiological implications. For instance, high energy states may represent less stable neural dynamics associated with schizophrenia since high energies correspond to low occurrence frequencies. Interestingly, the highest energy states within our data are ones featuring roughly equal numbers of super-threshold and sub-threshold regional activations, whereas the lowest energy states are homogeneous ones in which most regions showed super-threshold activation, or most regions showed sub-threshold activation (**Supplemental Figure S3**). Thus, high energy states may be those with the least coordination or most independence among regional activations, and the prevalence of high-energy states in schizophrenia may correspond to a lower degree of coordination across brain regions. Functionally, since the activation within the DMN is suppressed while engaging in a task (49), it is possible that overabundance of higher energy states i.e., more variability in the spontaneous activity in a clinical group, makes it more difficult to engage activation sequences associated with normal functioning. However, significantly lower energy states in bipolar disorder compared to controls suggest that the neural dynamics in bipolar disorder may be “overly stable” or spend more time in various attractors. Theoretically this may result in a hindered ability to make the brief excursions needed to engage in tasks. It is possible that patterns of recruitment of DMN regions in bipolar disorder (abnormal recruitment of dorsal DMN regions) and in schizophrenia (abnormal preferential recruitment of ventral DMN regions) observed in prior studies (50) contribute to such differences in energy distributions. Although major depression showed no difference in energy states compared to controls in the bilateral and left hemisphere DMN in our study, the presence of high energy states only in the right hemisphere DMN may suggest that a laterality of abnormalities in DMN circuitry could be pathophysiologically important. Although reduced DMN connectivity has been observed in major depression (51), laterality in connectional differences were not reported. Further studies are required to fully understand these differences and their clinical implications.

Pathophysiological interpretations based on the theoretical underpinnings of MEM can be proposed. Since fMRI signals localized to the gray matter are proxy to the underlying neural activity (6, 52, 53), it is plausible that fMRI data have Ising-like properties driven by neural connectivity and that some image-derived coefficients may relate to the MEM parameters, *J* and *h*. Previous studies (8) have reported that the *J* matrix from the pairwise MEM (**Equation 1**) explains the system interaction better than traditional FC and that strong connections in *J* are very likely to co-occur with diffusion streamline connections in the diffusion-MRI derived structural connectome (8). Other recent studies have examined the relationship between *J* and the structural connectome in more detail (7, 41).

We identified few attempts in the literature to find image-derived coefficients related to the first-order activation parameter, *h* (7). Activation rate, or average activation over time, is sometimes suggested, but for *z*-scored signal activation rates are fixed around 50%. In some contexts, particularly its native context of physics, the *h* parameter is described as the “external field.” This could be considered equivalent to the notion of the neural receptive field (54). This is not to suggest that the BOLD signal can be functionally localized and mapped to neural computation, rather that the *h* parameter in the MEM is theoretically related to the external environment. Since this study used resting fMRI there are no external stimuli source besides the fixation cross and the scanner noise. We suggest Equation 3 as the starting point for an image-derived representation of *h*.

Despite the strengths of the MEM studies discussed above, these models are encumbered by the exponential relationship between the number of unknown parameters and number of network nodes. Because of the exponential increase in the number of states that occurs with each additional region included, we consider a relatively small number of regions in defining states of the model. To mitigate this limitation, we have considered three different sub-systems. A further limitation of this approach is that it cannot be used to assess the interactions between the sub-systems directly; however, there is nonetheless relevance to studying smaller, localized systems within the larger context of the brain. This strategy represents a practical and reasonable option when guided by data-driven, hypothesis-based, or other well-delineated networks (as was the case here) to select nodes. Additionally, recent studies show that findings reported in second-order statistical FC studies (55) and first-order brain-wide association findings (56) are not very reliable. While we find that the MEM exhibits a strong relation to FC and can be used to estimate first-order nodal activity, more research is needed to examine the extent of reproducibility using the MEM. In our study, the findings were consistent between the bilateral and single hemisphere system for schizophrenia and bipolar disorder. In the case of major depression, it is difficult to say whether the right hemisphere finding is truly related to depression or an indication of results not being fully reproducible. Our schizophrenia/schizoaffective disorder sample was relatively small, the findings related to this group should be viewed with caution. However, our study on a slightly larger sample of adolescent-onset schizophrenia on a task fMRI data showed similar findings as well as showing correlation with executive function and negative symptoms (15), partly allaying the concerns of reproducibility.

We show that individual fMRI scans are amenable to MEM fitting techniques. Despite having fewer timepoints (490) than the number of nodes (512), we obtained high goodness-of-fit measures **(Fig 2)** due to the fact that many states had very low empirical probability, thus estimating their probability to be zero is not unreasonable. A fitness “sweep” that we conducted confirmed such a high-quality fit. To our knowledge, this is the first application of individual-level MEM to psychiatric neuroimaging. While the findings need to be replicated in independent samples, our results suggest that it may be possible to characterize clinical biomarkers using MEM-defined energy. In schizophrenia/schizoaffective disorder, DMN states were of higher energy than in controls, which is similar to our earlier finding on MEM-based examination of executive function task-fMRI data in adolescent-onset schizophrenia (15). The bipolar group had overall lower energy states than controls suggesting that the DMN may have a larger number of highly stable alternative resting states or spend more time in such states. For the depression group there may be abnormal DMN functional lateralization.

The translational goals of traditional FC studies (57), such as treatment response prediction (58), therapeutic targeting, biomarker identification (59, 60), and investigating pathophysiology can be tested within the MEM framework, which provides a potentially useful representation of the underlying biological connectivity. With advances in MRI hardware, it is now possible to acquire fMRI data at faster rates (shorter TRs) that could allow for even better estimation of *h* and *J* parameters in the MEM model. Even with more data, however, the correlations between predicted and observed probabilities of state occurrence will not be perfect because the MEM captures only the pairwise interactions among the nodes, not the higher order interactions, which are also not considered in traditional functional connectivity studies. The strengths of MEM paradigm are that it provides a principled, unified framework to derive first- and second-order model parameters associated with a theoretical system of interactions that yields accurate state probability predictions. Future research should focus on clinical translatability of this model and examine task-associated brain networks in relation to cognitive impairments and psychopathology.

## Data availability

The data is available through the UK Biobank data repository.

## Code availability

Implementations of this process, as well as equations and related MATLAB, R, and Python code can be found on GitHub (https://github.com/theisn/IsingModel_fMRI).

## Author Contributions

NT conceptualized the study analysis, performed analysis and drafted the manuscript; JB helped with MEM implementation, performed analysis and participated in editing the manuscript; JR supervised the analysis and participated in drafting and editing manuscript; JC and SI participated in the discussion of the analysis and edited the manuscript; KP managed the data procurement for the study, conceptualized the study design, supervised the analysis, and participated in drafting and editing manuscript.

## Supplemental Information

The regions included in the three models are shown visually in the **Supplemental Figure S1**.

**Supplemental Figure S1.**
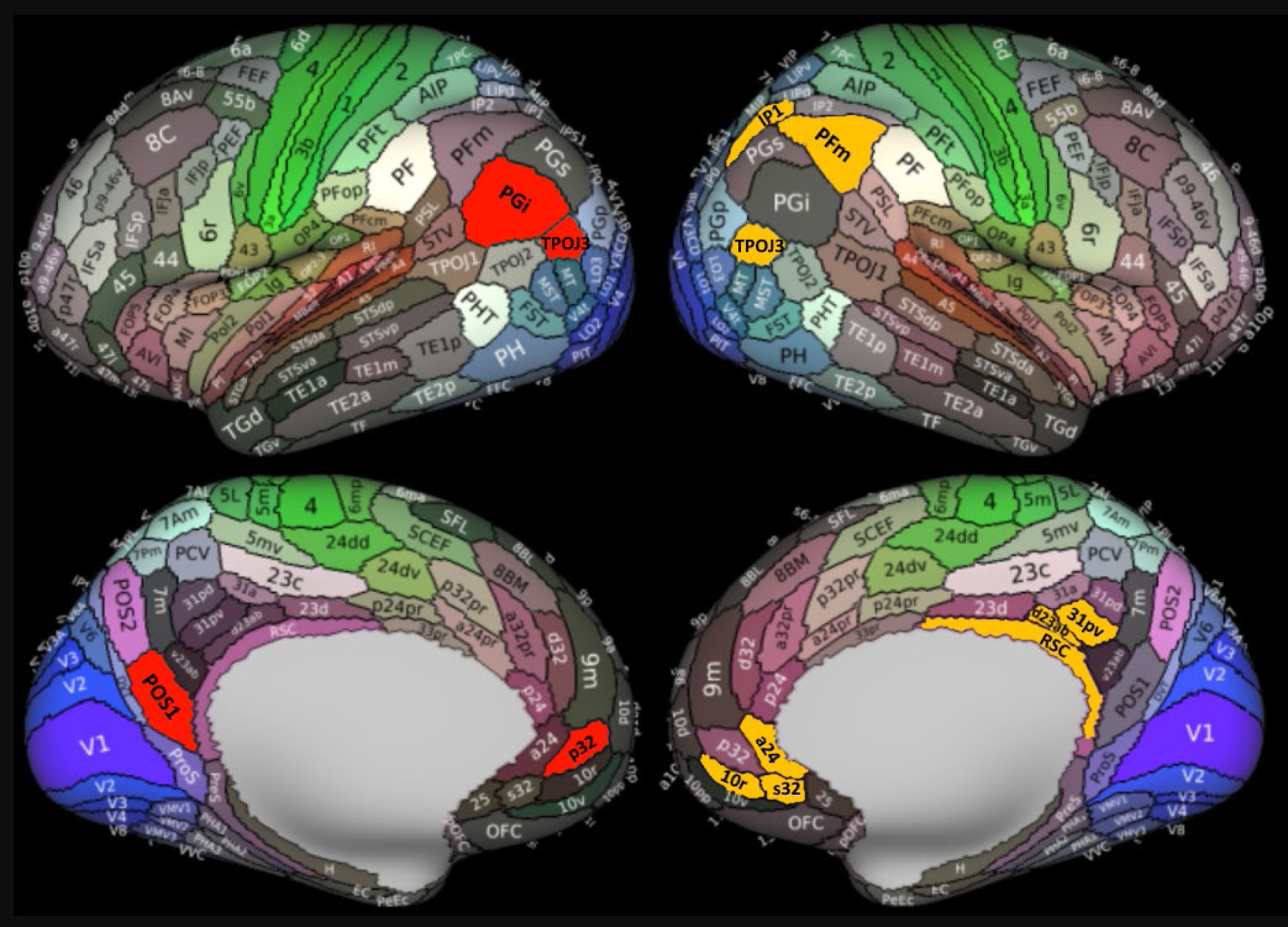
Brain Region Locations. Bright red ROIs represent the on the left indicate the bilateral system. Bright yellow ROIs on the right indicate both unilateral systems. This figure is a modified version of Figure 3 from Glasser et al. 2016, for a full explanation of label names outside the scope of this manuscript, as well as the meaning of the color map and other information, see reference (22).

### Supplemental Information 2

Substituting FC for *J* and equation (3) for *h* results in non-significant model fits (**Supplemental Figure S2**). Since the parameters in the Ising model (**Equation 1**) are unknown and must be computed using the MEM, and because the computed parameters from the MEM correlate significantly with image-derived variables (**Figure 3**), it is reasonable to assume that substituting the image-derived values for the MEM-derived values will also result in good fits, similar to those shown in (**Figure 2**). This does not turn out to be the case. Image-derived parameters result in poorer predictions of observed state occurrence probability.

**Supplemental Figure S2.**
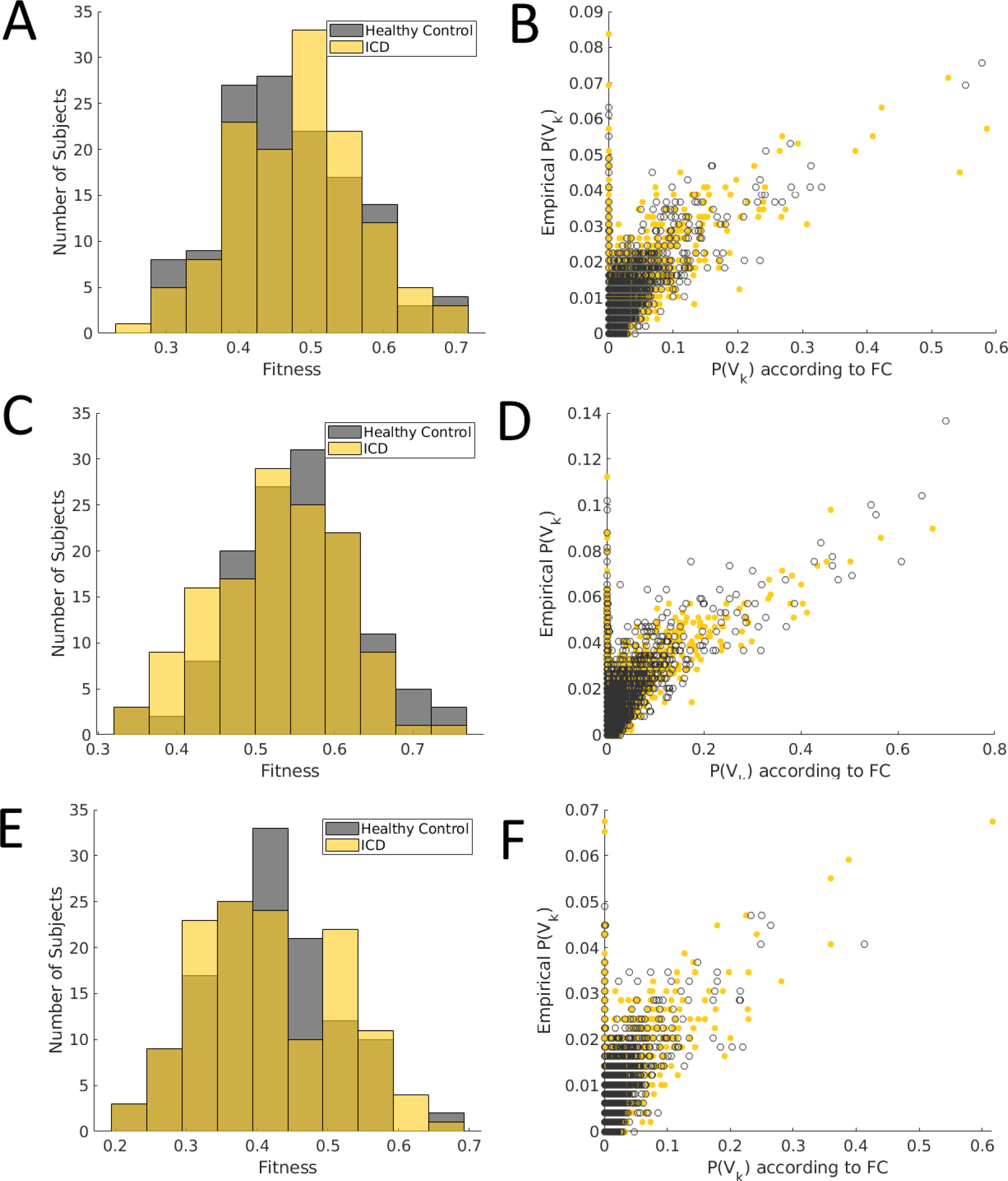
Fits according to image-derived coefficients. **Bilateral DMN nodes:** top row. **Left Hemisphere DMN nodes:** middle row. **Right Hemisphere DMN nodes:** bottom row. **A) bilateral:** two histograms, pooled ICD (N=132) in yellow and pooled HC (N=132) in black are compared. Fit (x-axis) is the correlation between predicted P(V_k_) according to the image-derived variables and observed *P(V_k_)* per individual. **B) bilateral:** all unique possible states are shown for all subjects according to their predicted probability (x-axis) versus their observed probability (y-axis). Black circles represent data from controls, yellow points represent data from the ICD group. **C) left hemisphere:** fit histograms for pooled ICD (yellow) and pooled HC (black) are compared. **D) left hemisphere:** all unique possible states are shown for all subjects according to their predicted probability (x-axis) versus their observed probability (y-axis). Black circles represent data from controls, yellow points represent data from the ICD group. **E) right hemisphere:** fit histograms for pooled ICD (yellow) and pooled HC (black) are compared. **F) right hemisphere:** all unique possible states are shown for all subjects according to their predicted probability (x-axis) versus their observed probability (y-axis). Black circles represent data from controls, yellow points represent data from the ICD group.

### Supplemental Information 3

Investigations into the relationships between MEM energy, functional connectivity, and first order binary activity are examined in (**Supplemental Figure S2**) for the bilateral, left hemisphere, and right hemisphere systems. Each subplot, **A-F**, depicts either the top 10% or bottom 10% of states by their energy, ranked according to the control group. The top chart in each subplot depicts the energy according to the state’s rank, in either decreasing or increasing order, depending if the bottom or top 10% are under consideration. The middle chart shows the total edge weight in the FC across all pairs of nodes that are both “on” for the state for the relevant 10% of states. The bottom chart illustrates the first order binary activity represented by the state.

**Supplemental Figure S3 A.**
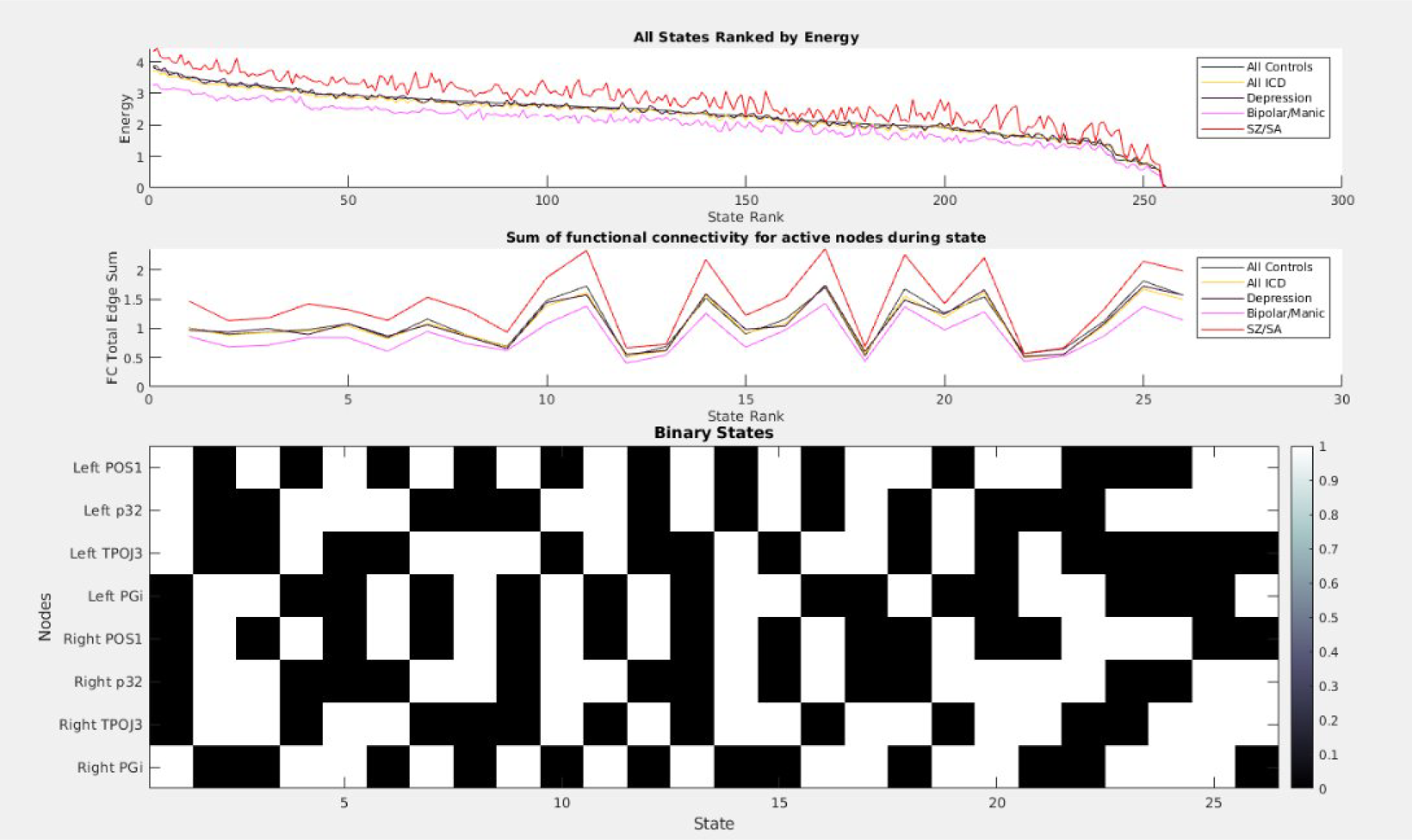
The top 10% of states in the bilateral system. Top chart: all states ranked by MEM energy of the control group. Middle chart: the second order co-activation of the top 10% of states, expressed as the sum of all edges between two “on” nodes in the FC. Bottom chart: the binary state configurations for the relevant states.

**Supplemental Figure S3 B.**
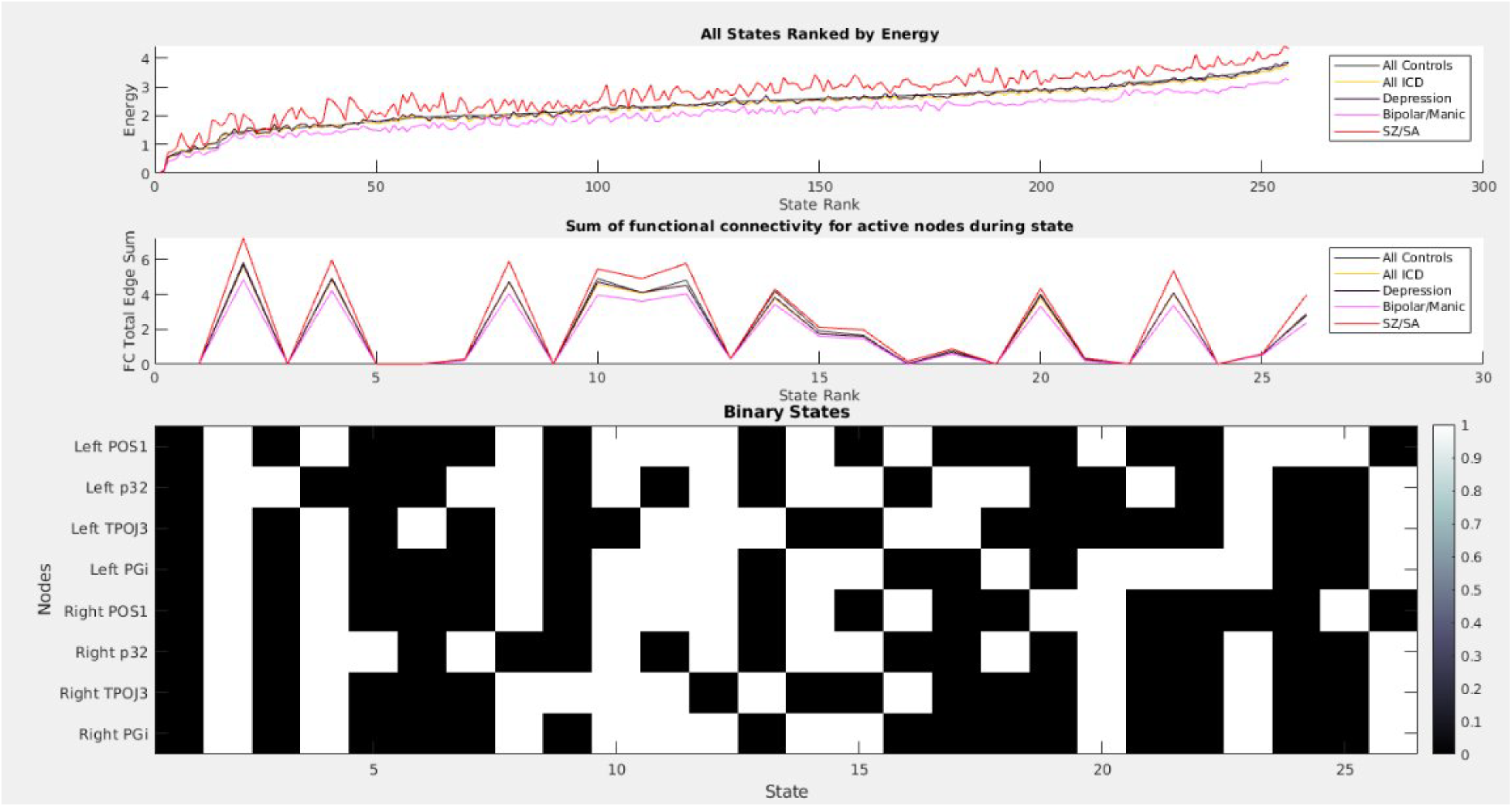
The bottom 10% of states in the bilateral system. Top chart: all states ranked by MEM energy of the control group. Middle chart: the second order co-activation of the bottom 10% of states, expressed as the sum of all edges between two “on” nodes in the FC. Bottom chart: the binary state configurations for the relevant states.

**Supplemental Figure S3 C.**
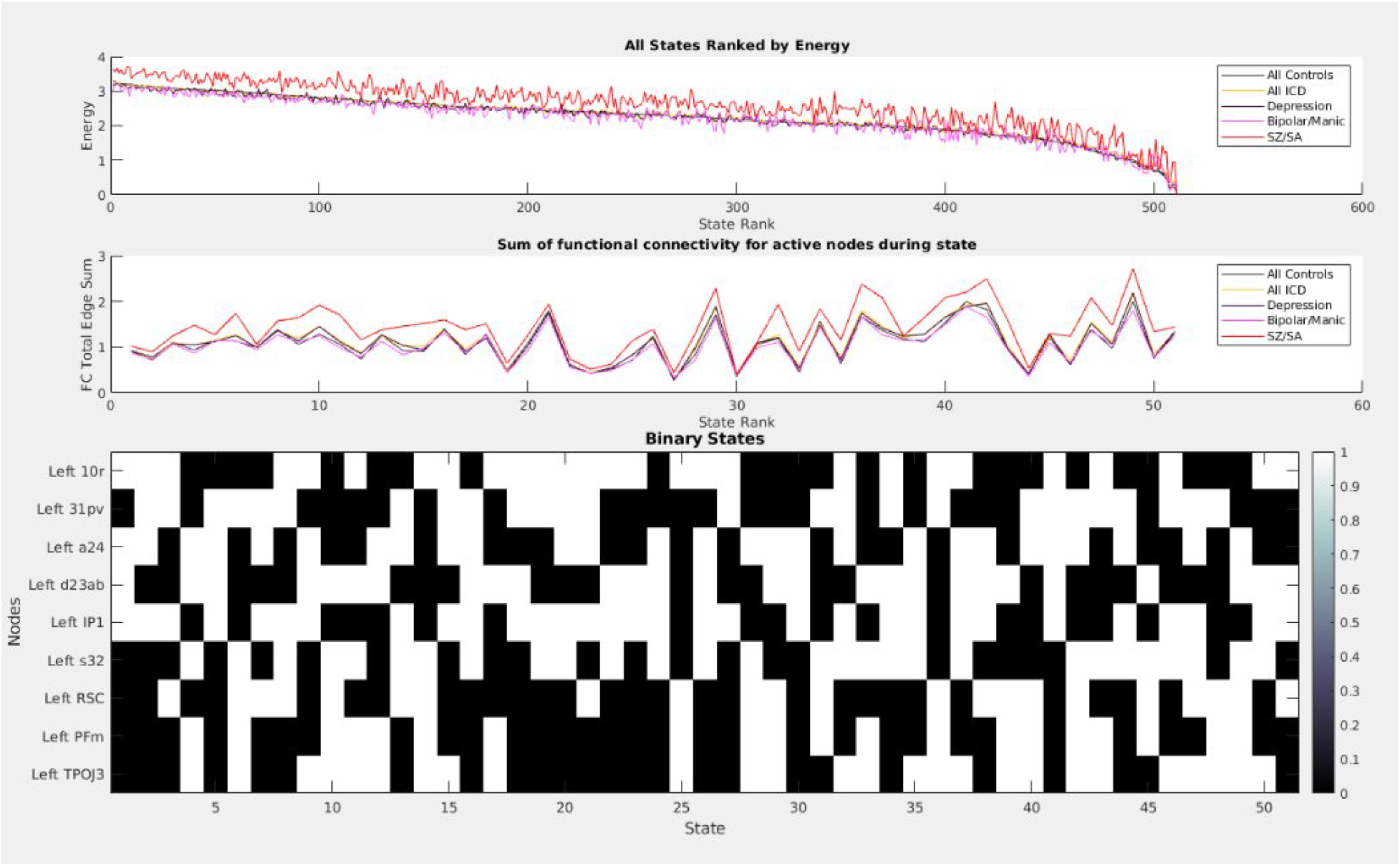
The top 10% of states in the left hemisphere system. Top chart: all states ranked by MEM energy of the control group. Middle chart: the second order co-activation of the top 10% of states, expressed as the sum of all edges between two “on” nodes in the FC. Bottom chart: the binary state configurations for the relevant states.

**Supplemental Figure S3 D.**
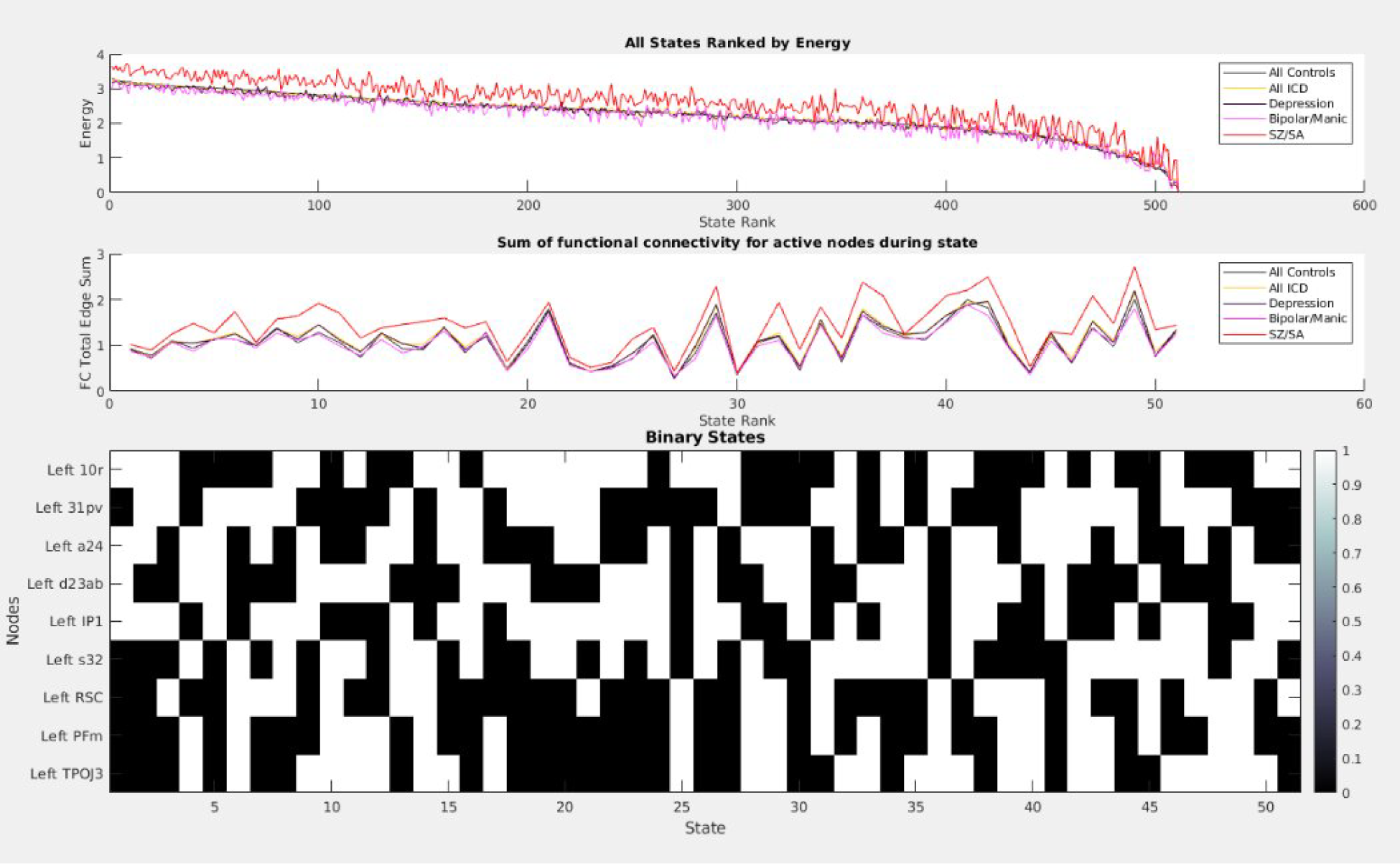
The bottom 10% of states in the left hemisphere system. Top chart: all states ranked by MEM energy of the control group. Middle chart: the second order co-activation of the bottom 10% of states, expressed as the sum of all edges between two “on” nodes in the FC. Bottom chart: the binary state configurations for the relevant states.

**Supplemental Figure S3 E.**
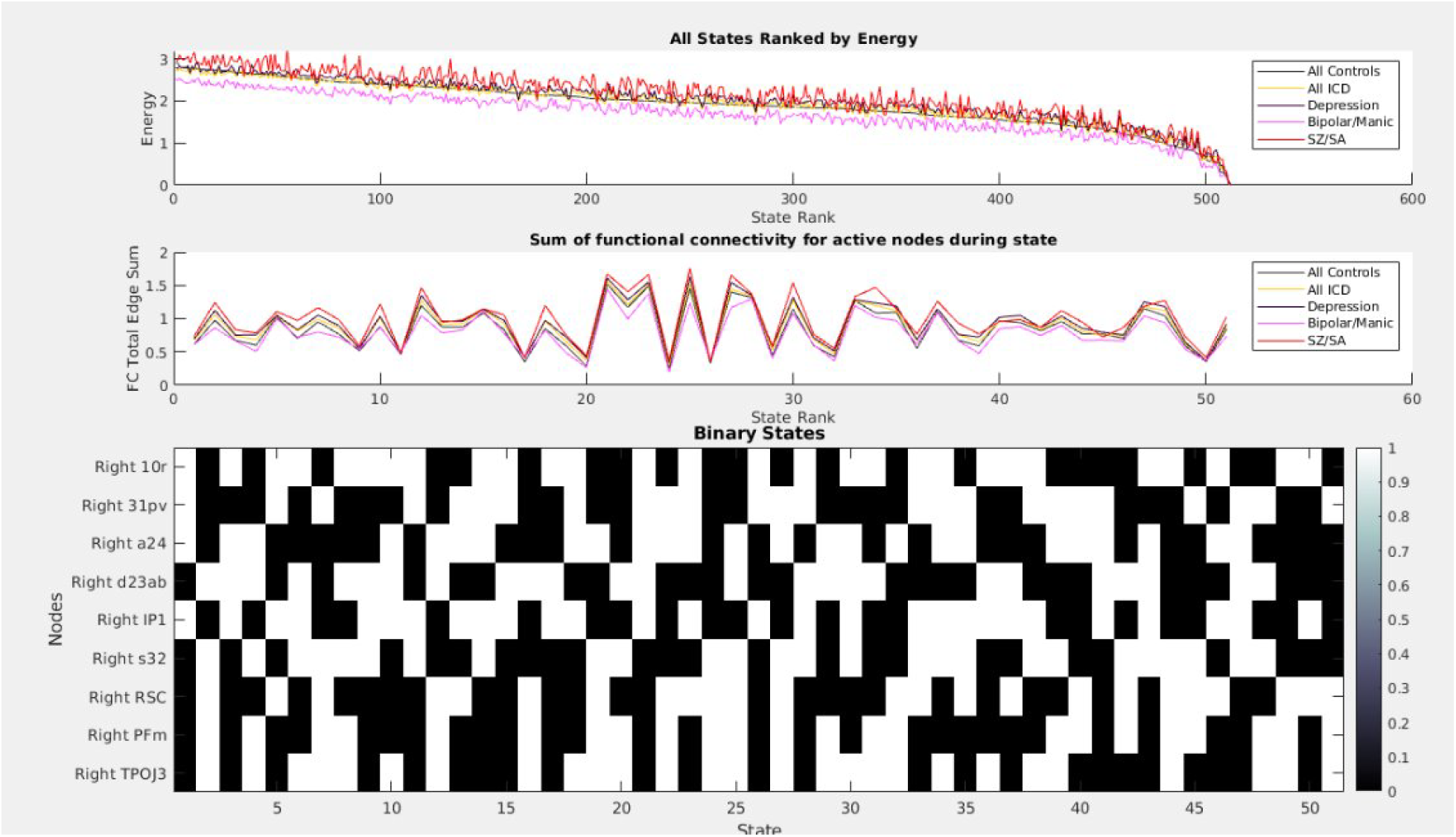
The top 10% of states in the right hemisphere system. Top chart: all states ranked by MEM energy of the control group. Middle chart: the second order co-activation of the top 10% of states, expressed as the sum of all edges between two “on” nodes in the FC. Bottom chart: the binary state configurations for the relevant states.

**Supplemental Figure S3 F.**
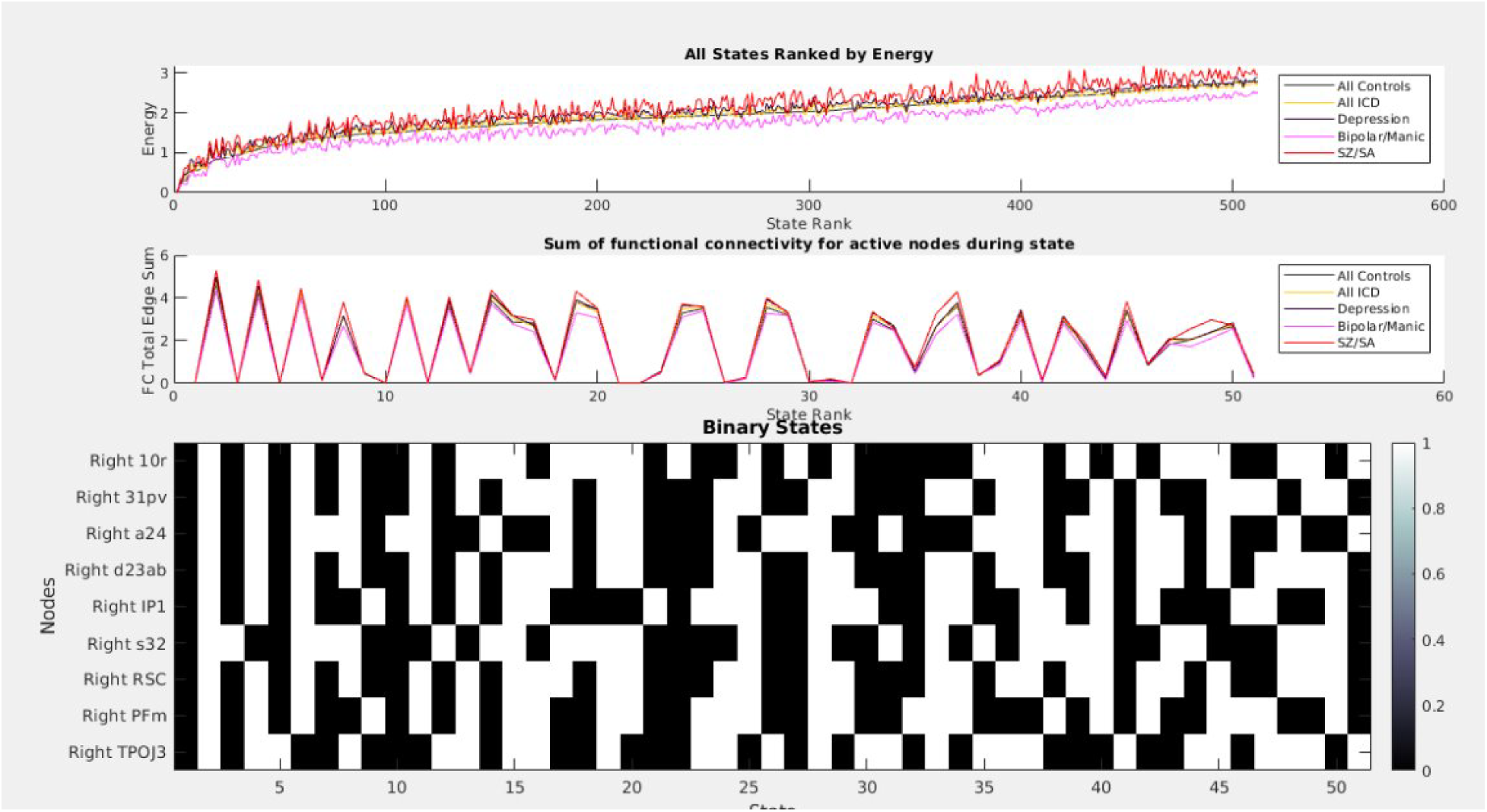
The bottom 10% of states in the right hemisphere system. Top chart: all states ranked by MEM energy of the control group. Middle chart: the second order co-activation of the bottom 10% of states, expressed as the sum of all edges between two “on” nodes in the FC. Bottom chart: the binary state configurations for the relevant states.

## REFERENCES

1. Amit DJ, Gutfreund H, Sompolinsky H (1985): Spin-glass models of neural networks. Phys Rev A Gen Phys. 32:1007–1018.

2. Fraiman D, Balenzuela P, Foss J, Chialvo DR (2009): Ising-like dynamics in large-scale functional brain networks. Phys Rev E Stat Nonlin Soft Matter Phys. 79:061922.

3. Hopfield JJ (1982): Neural networks and physical systems with emergent collective computational abilities. Proceedings of the National Academy of Sciences of the United States of America. 79:2554–2558.

4. Chialvo DR (2010): Emergent complex neural dynamics. Nature Physics. 6:744–750.

5. Gilson M, Moreno-Bote R, Ponce-Alvarez A, Ritter P, Deco G (2016): Estimation of Directed Effective Connectivity from fMRI Functional Connectivity Hints at Asymmetries of Cortical Connectome. PLoS Comput Biol. 12:e1004762.

6. Drew PJ (2019): Vascular and neural basis of the BOLD signal. Curr Opin Neurobiol. 58:61–69.

7. Kang J, Jeong SO, Pae C, Park HJ (2021): Bayesian estimation of maximum entropy model for individualized energy landscape analysis of brain state dynamics. Human brain mapping. 42:3411–3428.

8. Watanabe T, Hirose S, Wada H, Imai Y, Machida T, Shirouzu I, et al. (2013): A pairwise maximum entropy model accurately describes resting-state human brain networks. Nat Commun. 4:1370.

9. Sharma A, Wolf DH, Ciric R, Kable JW, Moore TM, Vandekar SN, et al. (2017): Common Dimensional Reward Deficits Across Mood and Psychotic Disorders: A Connectome-Wide Association Study. The American journal of psychiatry. 174:657–666.

10. Das TK, Abeyasinghe PM, Crone JS, Sosnowski A, Laureys S, Owen AM, Soddu A (2014): Highlighting the structure-function relationship of the brain with the Ising model and graph theory. Biomed Res Int. 2014:237898.

11. Yeh FC, Tang AN, Hobbs JP, Hottowy P, Dabrowski W, Sher A, et al. (2010): Maximum Entropy Approaches to Living Neural Networks. Entropy. 12:89–106.

12. Watanabe T, Hirose S, Wada H, Imai Y, Machida T, Shirouzu I, et al. (2014): Energy landscapes of resting-state brain networks. Frontiers in neuroinformatics. 8:12.

13. Ezaki T, Watanabe T, Ohzeki M, Masuda N (2017): Energy landscape analysis of neuroimaging data. Philos Trans A Math Phys Eng Sci. 375:20160287.

14. Gu S, Cieslak M, Baird B, Muldoon SF, Grafton ST, Pasqualetti F, Bassett DS (2018): The Energy Landscape of Neurophysiological Activity Implicit in Brain Network Structure. Sci Rep. 8:2507.

15. Theis N, Bahuguna J, Rubin J, Muldoon B, Prasad KM (2023): Energy in functional brain states correlates with cognition in adolescent schizophrenia and healthy persons. bioRxiv.

16. Jeong SO, Kang J, Pae C, Eo J, Park SM, Son J, Park HJ (2021): Empirical Bayes estimation of pairwise maximum entropy model for nonlinear brain state dynamics. NeuroImage. 244:118618.

17. Piantadosi S, Byar DP, Green SB (1988): The ecological fallacy. Am J Epidemiol. 127:893–904.

18. Birn RM, Molloy EK, Patriat R, Parker T, Meier TB, Kirk GR, et al. (2013): The effect of scan length on the reliability of resting-state fMRI connectivity estimates. NeuroImage. 83:550–558.

19. Ball TM, Goldstein-Piekarski AN, Gatt JM, Williams LM (2017): Quantifying person-level brain network functioning to facilitate clinical translation. Translational psychiatry. 7:e1248.

20. Gratton C, Kraus BT, Greene DJ, Gordon EM, Laumann TO, Nelson SM, et al. (2020): Defining Individual-Specific Functional Neuroanatomy for Precision Psychiatry. Biological psychiatry. 88:28–39.

21. Littlejohns TJ, Holliday J, Gibson LM, Garratt S, Oesingmann N, Alfaro-Almagro F, et al. (2020): The UK Biobank imaging enhancement of 100,000 participants: rationale, data collection, management and future directions. Nat Commun. 11:2624.

22. Glasser MF, Coalson TS, Robinson EC, Hacker CD, Harwell J, Yacoub E, et al. (2016): A multi-modal parcellation of human cerebral cortex. Nature. 536:171–178.

23. Sandhu Z, Tanglay O, Young IM, Briggs RG, Bai MY, Larsen ML, et al. (2021): Parcellation-based anatomic modeling of the default mode network. Brain Behav. 11:e01976.

24. Garrity AG, Pearlson GD, McKiernan K, Lloyd D, Kiehl KA, Calhoun VD (2007): Aberrant “default mode” functional connectivity in schizophrenia. The American journal of psychiatry. 164:450–457.

25. Broyd SJ, Demanuele C, Debener S, Helps SK, James CJ, Sonuga-Barke EJ (2009): Default-mode brain dysfunction in mental disorders: a systematic review. Neurosci Biobehav Rev. 33:279–296.

26. Whitfield-Gabrieli S, Ford JM (2012): Default mode network activity and connectivity in psychopathology. Annu Rev Clin Psychol. 8:49–76.

27. Doucet GE, Bassett DS, Yao N, Glahn DC, Frangou S (2017): The Role of Intrinsic Brain Functional Connectivity in Vulnerability and Resilience to Bipolar Disorder. The American journal of psychiatry. 174:1214–1222.

28. Doucet GE, Janiri D, Howard R, O’Brien M, Andrews-Hanna JR, Frangou S (2020): Transdiagnostic and disease-specific abnormalities in the default-mode network hubs in psychiatric disorders: A meta-analysis of resting-state functional imaging studies. European psychiatry : the journal of the Association of European Psychiatrists. 63:e57.

29. Miller KL, Alfaro-Almagro F, Bangerter NK, Thomas DL, Yacoub E, Xu J, et al. (2016): Multimodal population brain imaging in the UK Biobank prospective epidemiological study. Nature neuroscience. 19:1523–1536.

30. Smith S, Alfaro-Almagro F, Miller K (2020): UK Biobank Brain Imaging Documentation. Wellcome Centre for Integrative Neuroimaging (WIN-FMRIB), Oxford University on behalf of UK Biobank: http://www.ukbiobank.ac.uk.

31. Jenkinson M, Beckmann CF, Behrens TE, Woolrich MW, Smith SM (2012): Fsl. NeuroImage. 62:782–790.

32. Beckmann CF, Smith SM (2004): Probabilistic independent component analysis for functional magnetic resonance imaging. IEEE transactions on medical imaging. 23:137–152.

33. Salimi-Khorshidi G, Douaud G, Beckmann CF, Glasser MF, Griffanti L, Smith SM (2014): Automatic denoising of functional MRI data: combining independent component analysis and hierarchical fusion of classifiers. NeuroImage. 90:449–468.

34. Buitrago P, Nystrom N (2021): Neocortex and Bridges-2: A High Performance AI+HPC Ecosystem for Science, Discovery, and Societal Good High Performance Computing, pp 205–219.

35. Neurolab C (2017): HCP-MMP1.0 volumetric (NIfTI) masks in native structural space. figshare, pp Dataset.

36. Mills K (2016): HCP-MMP1.0 projected on fsaverage. In: figshare, editor.

37. Fischl B (2012): FreeSurfer. NeuroImage. 62:774–781.

38. Kloucek MB, Machon T, Kajimura S, Royall CP, Masuda N, Turci F (2023): Biases in inverse Ising estimates of near-critical behavior. Phys Rev E. 108:014109.

39. Ising E (1924): Beitrag zur Theorie des Ferro- und Paramagnetismus: University of Hamburg.

40. Ising T, Folk R, Kenna R, Berche B, Holovatch Y (2017): The Fate of Ernst Ising and the Fate of his Model. arXiv: History and Philosophy of Physics.

41. Ashourvan A, Shah P, Pines A, Gu S, Lynn CW, Bassett DS, et al. (2021): Pairwise maximum entropy model explains the role of white matter structure in shaping emergent co-activation states. Commun Biol. 4:210.

42. Knuuttila T, Loettgers A (2014): Magnets, Spins, and Neurons: The Dissemination of Model Templates across Disciplines. Monist. 97:280–300.

43. Jo Y, Faskowitz J, Esfahlani FZ, Sporns O, Betzel RF (2021): Subject identification using edge-centric functional connectivity. NeuroImage. 238:118204.

44. Liu TT, Nalci A, Falahpour M (2017): The global signal in fMRI: Nuisance or Information? NeuroImage. 150:213–229.

45. Jaynes ET (1957): Information Theory and Statistical Mechanics. Physical Review. 106:620–630.

46. Huang CC, Luo Q, Palaniyappan L, Yang AC, Hung CC, Chou KH, et al. (2020): Transdiagnostic and Illness-Specific Functional Dysconnectivity Across Schizophrenia, Bipolar Disorder, and Major Depressive Disorder. Biol Psychiatry Cogn Neurosci Neuroimaging. 5:542–553.

47. Brandl F, Avram M, Weise B, Shang J, Simoes B, Bertram T, et al. (2019): Specific Substantial Dysconnectivity in Schizophrenia: A Transdiagnostic Multimodal Meta-analysis of Resting-State Functional and Structural Magnetic Resonance Imaging Studies. Biological psychiatry. 85:573–583.

48. Li C, Dong M, Womer FY, Han S, Yin Y, Jiang X, et al. (2021): Transdiagnostic time-varying dysconnectivity across major psychiatric disorders. Human brain mapping. 42:1182–1196.

49. Raichle ME (2015): The brain’s default mode network. Annu Rev Neurosci. 38:433–447.

50. Ongur D, Lundy M, Greenhouse I, Shinn AK, Menon V, Cohen BM, Renshaw PF (2010): Default mode network abnormalities in bipolar disorder and schizophrenia. Psychiatry research. 183:59–68.

51. Yan CG, Chen X, Li L, Castellanos FX, Bai TJ, Bo QJ, et al. (2019): Reduced default mode network functional connectivity in patients with recurrent major depressive disorder. Proceedings of the National Academy of Sciences of the United States of America. 116:9078–9083.

52. Logothetis NK (2007): The ins and outs of fMRI signals. Nature neuroscience. 10:1230–1232.

53. Hillman EM (2014): Coupling mechanism and significance of the BOLD signal: a status report. Annu Rev Neurosci. 37:161–181.

54. Spillmann L, Dresp-Langley B, Tseng CH (2015): Beyond the classical receptive field: The effect of contextual stimuli. J Vis. 15:7.

55. Noble S, Scheinost D, Constable RT (2019): A decade of test-retest reliability of functional connectivity: A systematic review and meta-analysis. NeuroImage. 203:116157.

56. Marek S, Tervo-Clemmens B, Calabro FJ, Montez DF, Kay BP, Hatoum AS, et al. (2022): Reproducible brain-wide association studies require thousands of individuals. Nature. 603:654–660.

57. van den Heuvel MP, Hulshoff Pol HE (2010): Exploring the brain network: a review on resting-state fMRI functional connectivity. Eur Neuropsychopharmacol. 20:519–534.

58. Goff DC, Roffman J, Holt DJ (2023): Another Step Toward the Prediction of Antipsychotic Treatment Response Using Functional Connectivity. The American journal of psychiatry. 180:787–788.

59. Kraguljac NV, McDonald WM, Widge AS, Rodriguez CI, Tohen M, Nemeroff CB (2021): Neuroimaging Biomarkers in Schizophrenia. The American journal of psychiatry. 178:509–521.

60. Keshavan MS, Collin G, Guimond S, Kelly S, Prasad KM, Lizano P (2020): Neuroimaging in Schizophrenia. Neuroimaging Clin N Am. 30:73–83.

